# Network controllability analysis reveals the antiviral potential of Etravirine against Hepatitis E Virus infection

**DOI:** 10.1101/2024.06.21.600064

**Authors:** Shabnam Ansari, Dipanka Tanu Sarmah, Rohit Verma, Kannan Chandrasekar, Samrat Chatterjee, Milan Surjit

**Affiliations:** Virology Laboratory, Centre for Virus research, therapeutics and vaccines, Translational Health Science and Technology Institute, NCR Biotech Science Cluster, Faridabad, Haryana, India; Complex analysis group, Computational and Mathematical Biology Centre, Translational Health Science and Technology Institute, NCR Biotech Science Cluster, Faridabad, Haryana, India

**Keywords:** Hepatitis E Virus, Connectivity Map, Network controllability, Casein Kinase 1ε, Protein kinase B, Etravirine

## Abstract

Hepatitis E virus (HEV) is a major cause of acute viral hepatitis in lower- and middle-income countries. HEV infection may lead to acute liver failure, chronic liver disease and high mortality in pregnant women. Antiviral therapy is not a standard treatment for HEV patients. Computational biology tools promise to revolutionize the antiviral drug discovery. Here, we analyzed the transcriptome data of HEV infected primary human hepatocyte (PHH)-cells through connectivity map database and applied control theory on functional network to identify antiviral targets against HEV. The above analyses predicted PKCβ, PKB/AKT and CK1ε as potential antiviral targets against HEV. The antiviral function of PKB/AKT and CK1ε was experimentally validated by using respective biochemical inhibitors in g3 (genotype 3)-HEV replicon and Huh7 cell-based model of g3 and g1-HEV infection. Further, knockdown of CK1ε showed a similar effect. These data confirmed that CK1ε is an antiviral target for HEV. At present, there are no FDA approved drugs targeting CK1ε. Etravirine is an FDA approved non-nucleoside reverse transcriptase inhibitor drug, used for the treatment of Human immunodeficiency virus type 1 (HIV-1) infected patients. An *in silico* study predicted Etravirine to be a potent inhibitor of CK1ε. Our experiments revealed potent antiviral activity of Etravirine against HEV, which was mediated via its ability to inhibit the activity of CK1ε. Taken together, the current study demonstrates that PKB/AKT and CK1ε are bonafide antiviral targets for HEV and paves the way for repurposing Etravirine for the treatment of HEV infected patients.

**Importance:** Antiviral treatment is not the standard care for acute viral hepatitis E patients. Unbiased identification of antiviral targets or large-scale screening of antiviral compounds against the hepatitis E virus (HEV) has not been reported. Here, computational biology approach was followed to unbiasedly identify antiviral targets of HEV. Transcriptome data of HEV infected primary human hepatocyte (PHH) cells were analyzed to identify modulators of the network and generate directional networks. Network controllability analysis identified PKCβ, PKB/AKT and CK1ε as potential antiviral targets against HEV. Antiviral function of PKB/AKT and CK1ε was confirmed using cell-based models of genotype 1 (g1)- and g3-HEV infection. Further experiments demonstrated the antiviral activity of Etravirine against HEV, mediated via its ability to inhibit the CK1ε activity. Etravirine is an FDA approved non-nucleoside reverse transcriptase inhibitor, used for the treatment of Human immunodeficiency virus type-1 (HIV-1)-infected patients. This study reveals the potential of repurposing Etravirine for treatment of HEV patients and illustrate the importance of computational biology in antiviral drug discovery.

## Introduction

Hepatitis E Virus (HEV) is a major cause of viral hepatitis globally. It causes acute viral hepatitis (AVH) in humans, characterized by jaundice, anorexia, nausea, abdominal pain, malaise, fever, and hepatomegaly, which may progress to acute liver failure (ALF). It also causes chronic hepatitis, fulminant hepatitis, and extrahepatic manifestations such as Guillain-Barre syndrome and neurological amyotrophy in a subset of patients. The disease worsens during pregnancy, with a 20-25% mortality rate. Recent reports suggest chronic HEV infection in ∼60% of liver transplantation patients (1–4).

Antiviral treatment is not the standard care for acute viral hepatitis E patients. Ribavirin monotherapy, pegylated interferon alpha or a combination of both is considered for viral clearance in chronic hepatitis E, immunocompromised and organ transplant patients (5–7). However, the side effects of both ribavirin and interferon therapy render the treatment unsuitable for several categories of patients (8–10).

Few studies have been undertaken to identify specific antivirals against HEV. Notably, studies on virus-targeting antiviral screening have identified 3-(4-Hydroxyphenyl) propionic acid [HPPA (viral methyltransferase inhibitor)], Favipiravir [viral RNA-dependent RNA polymerase (RdRp) inhibitor] in combination with Sofosbuvir (viral RdRp inhibitor), Methotrexate (viral helicase inhibitor), 2’-C-methylcytidine (viral RdRp inhibitor), 66E2 (viral RdRp inhibitor) as potential antivirals against HEV (11–15). Studies on host-targeting antivirals have identified compounds such as Mycophenolic acid, Artesunate and Fenofibrate as potential antivirals against HEV (16–18). Our earlier study identified anti-HEV activity of a cyclic peptide (CP11), which acts by interrupting the interaction between HEV ORF3 and host TSG101 (Tumor susceptibility gene 101), leading to the inhibition of progeny virus release (19). Our earlier study also demonstrated the antiviral potential of zinc salts and zinc Oxide nanoparticles against HEV (20, 21). Towards identifying potent antiviral targets against HEV in the host, virus-host protein-protein interaction (PPI) network has been explored by yeast two hybrid-based cDNA library screening approach or immunoprecipitation-mass spectrometry (IP-MS) approach using viral proteins as bait (22, 23). Although several potential targets were identified, functional validation of the targets and their small molecule modulators remain unexplored. In summary, although some progress has been made towards identifying specific antivirals against HEV following target specific approach or small-scale focused screening approach, unbiased identification of antiviral targets or large-scale screening of antiviral compounds against HEV has not been reported.

A major obstacle in performing high-throughput antiviral screening assays in HEV infected cells is attributed to the lack of an efficient cell-culture based infection model of the virus. Alternatively, computational biology methods may be explored to identify antiviral targets and antiviral drugs against HEV using transcriptome and/or proteome profile data of HEV infected cells and respective controls. One of the approaches for computational identification of antiviral targets is based on analysis of the differential gene expression dataset of a diseased/infected/perturbed sample in the connectivity map (CMap) database, construction of a directional protein-protein interaction (PPI) network, followed by controllability analysis of the network (24, 25). CMap is a reference drug perturbation dataset containing transcriptome profile of multiple cell lines treated with drugs. Comparison of transcriptome profile of a disease/infection sample and reference transcriptome profiles available in CMap helps in identifying the inverse drug-disease/infection relation. Control theory analysis of the resulting data helps identify the network’s most fragile nodes. Therefore, a potential target obtained through a combination of CMap and network controllability analysis will not only regulate the network dynamics, but also, likely induce a transition in the state of the system from “disease” to “healthy” (26).

Todt et al. have reported the host transcriptome profile in g3-HEV infected PHH (Primary human hepatocyte) cells (27). Raw data from the Gene Expression Omnibus (GEO) database was used to select differentially expressed genes (DEGs) at multiple cut offs and analyze them in the CMap database to identify modulators. Subsequently, DEGs and modulators were used to construct directional PPI networks, followed by network controllability analysis to identify the indispensable modulators. Further analysis identified PRKCB, AKT1 and CSNK1E as potential antiviral targets. Experimental evaluation of HEV inhibitory activity of AKT and CK1ε inhibitors revealed anti-HEV activity of respective inhibitor compounds. Further experiments confirmed the antiviral activity of Etravirine against HEV.

Etravirine is a diarylpyrimidine group drug used for the treatment of Human immunodeficiency virus type 1 (HIV-1) infected patients. It is a second-generation non-nucleoside reverse transcriptase inhibitor, with the ability to inhibit the HIV-1 strains that are resistant to many other anti-retroviral drugs (28). It has a half-life of around 41 hours and it gets metabolized in the liver by CYP3A4, CYP2C9 and CYP2C19 enzymes (29). The drug lacks any serious adverse effect except that it leads to development of rashes in some (30). Drug repurposing study has shown the therapeutic utility of Etravirine against Friedreich’s ataxia, a genetic disorder of progressive loss of coordinated muscles strength (31). In case of Friedreich’s ataxia, Etravirine acts by increasing the level of frataxin by promoting its translation (31). In a recent study, a search for casein kinase 1 epsilon (CK1ε) inhibitor from a database of FDA-approved drugs identified Etravirine as a potential candidate (32). CK1 family of protein kinases are important in multiple biological pathways. CK1ε shares around 96% sequence homology with CK1δ. Both kinases are regulated by phosphorylation of their C-terminal tails. In turn, they phosphorylate a set of substrates and control many biological processes, including circadian cycle [PER (period) 2 phosphorylation], cell-cell interaction and cancer (connexin-43 phosphorylation), Cap-dependent translation, regulation of AMPK (Adenosine monophosphate-activated protein kinase) pathway and cancer (4EBP1 phosphorylation), dopamine signaling and drug addiction [DARPP (Dopamine and cAMP-regulated phosphoprotein)-32 phosphorylation] (33). CK1ε also phosphorylates TRAF3 (Tumor necrosis factor receptor associated factor 3) and regulates the antiviral innate immune response (34). Given the importance of CK1ε in multiple cellular processes and during viral infection, the potential use of Etravirine as an antiviral drug against HEV is discussed.

## Results

### Transcriptome profile analysis of HEV infected PHH cells to identify antiviral targets

Raw data of transcriptome profile of 48hr g3-HEV infected primary human hepatocyte (PHH) cells and corresponding uninfected cells were analysed to identify the differentially expressed genes (DEGs) (Figure 1A). Note that Todt et al. reported a peak level of viral replication to occur at 48 hour post-infection and highest number of DEGs were observed at that time point (27). The initial transcriptome dataset comprised 21,475 genes, and subsequent data filtration reduced it to 15,094 genes. DEG analysis was performed at 1.5, 2 and 2.5 fold cut offs to ensure that our findings do not unduly rely on a single arbitrary criterion. It also allows detection of a wide range of biologically important variances, from minor (1.5-fold, hereafter denoted as C1.5) to moderate (2-fold, hereafter denoted as C2.0) and comparatively severe (2.5-fold, hereafter denoted as C2.5) variations. 1299, 653, 382 genes were upregulated and 1386, 515, 348 genes were downregulated at C1.5, C2 and C2.5, respectively (Figure 1B). Since the CMap portal allows only the top 150 up-and down-regulated genes, CMap analysis for each category resulted in the same set of 36 modulators. The top 20 Biological processes and pathways associated with these genes are shown (Figure 1C and 1D). Note that the list is dominated by serotonergic receptor encoding genes (11 gene), Casein kinase encoding genes (4 genes) and adenosine receptor encoding genes (4 genes) (Table S1).

**Figure 1.**
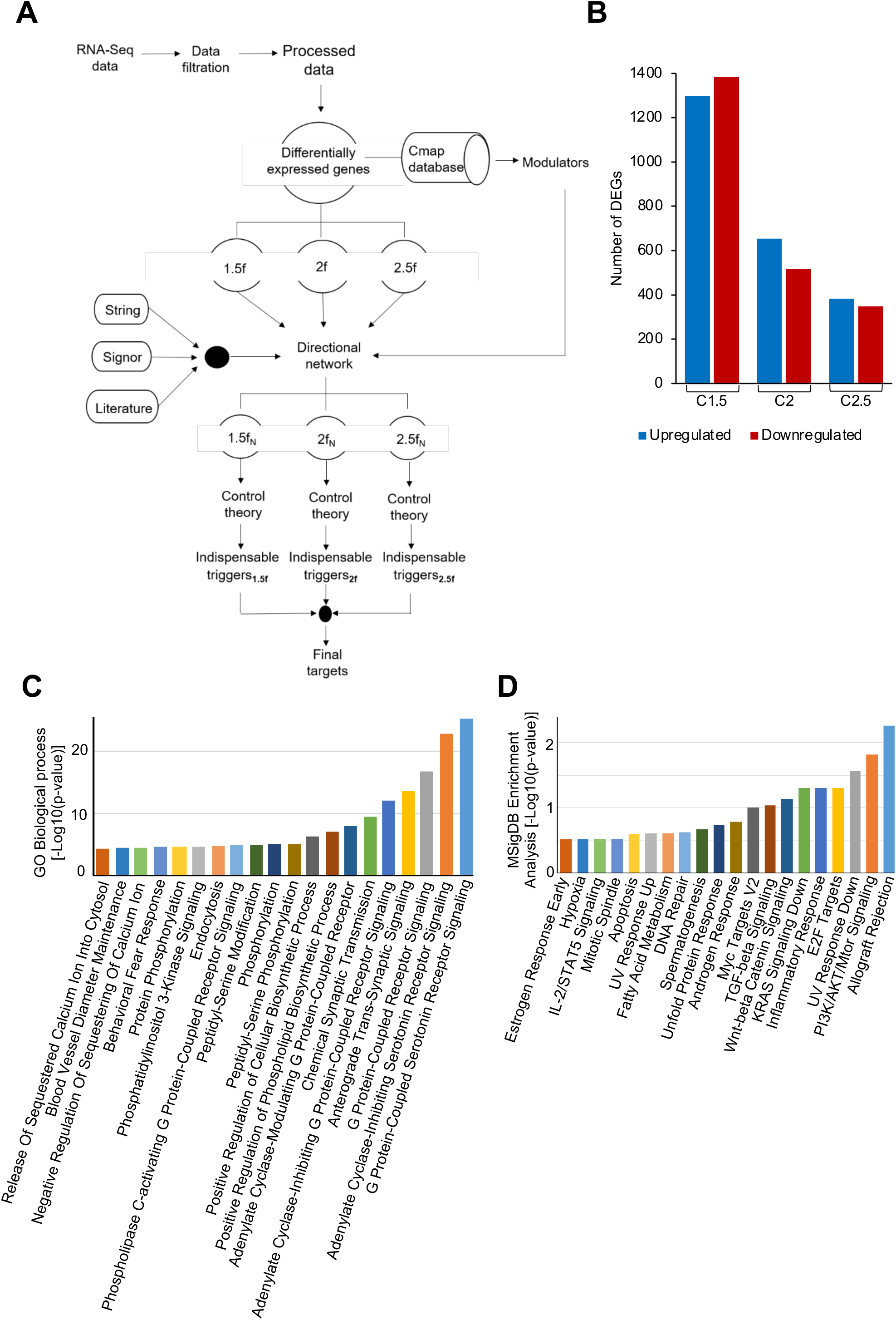
*In silico* analysis of transcriptome of g3-HEV infected PHH cells. A) Schematic of *in silico* analysis B) Number of DEGs identified at the indicated cut offs, X-axis: fold change; Y-axis, number of upregulated (Blue bar) and downregulated (red bar) genes. C) Top 20 “GO biological processes” associated with the modulators. D) Top 20 “pathways” associated with the modulators obtained by MSigDB enrichment analysis.

Next, a directional protein-protein interaction network (DPN) was generated by combining STRING (version 11.0), Signor (version 2.0) and literature and DEGs and modulators of each category were mapped to it (Figure 2A-2C). 1314, 600 and 373 nodes and 4398, 1741 and 1054 edges were detected at C1.5, C2 and C2.5 cut off, respectively (Figure 2D). Each displayed one giant component and many small disconnected components (Figure 2A-C). Analysis of the network properties revealed that C1.5 has the highest betweenness centrality (0.7992), indicating that nodes in C1.5 play a crucial role in connecting other nodes in the network. C2 stands next, followed by C2.5 (Figure 2E). C2 has the highest closeness centrality (0.2251), suggesting that, on average, nodes in C2 are closer to each other than the other two (Figure 2E). This easiness in traversing is also reflected in the fact that it has the shortest average path length (2.4269) (Figure 2E). C2.5 has the highest clustering coefficient (0.0883), indicating a higher degree of interconnectedness among neighbours, compared to C1.5 and C2. C1.5 has the highest indegree and outdegree, suggesting that nodes in C1.5 are more connected and receive/send more edges than the other subnetworks. It also has the highest neighbourhood degree (23.0393), indicating a higher density of connections in the immediate vicinity of nodes. The values of the centrality measures for each node are provided in Table S2.

**Figure 2.**
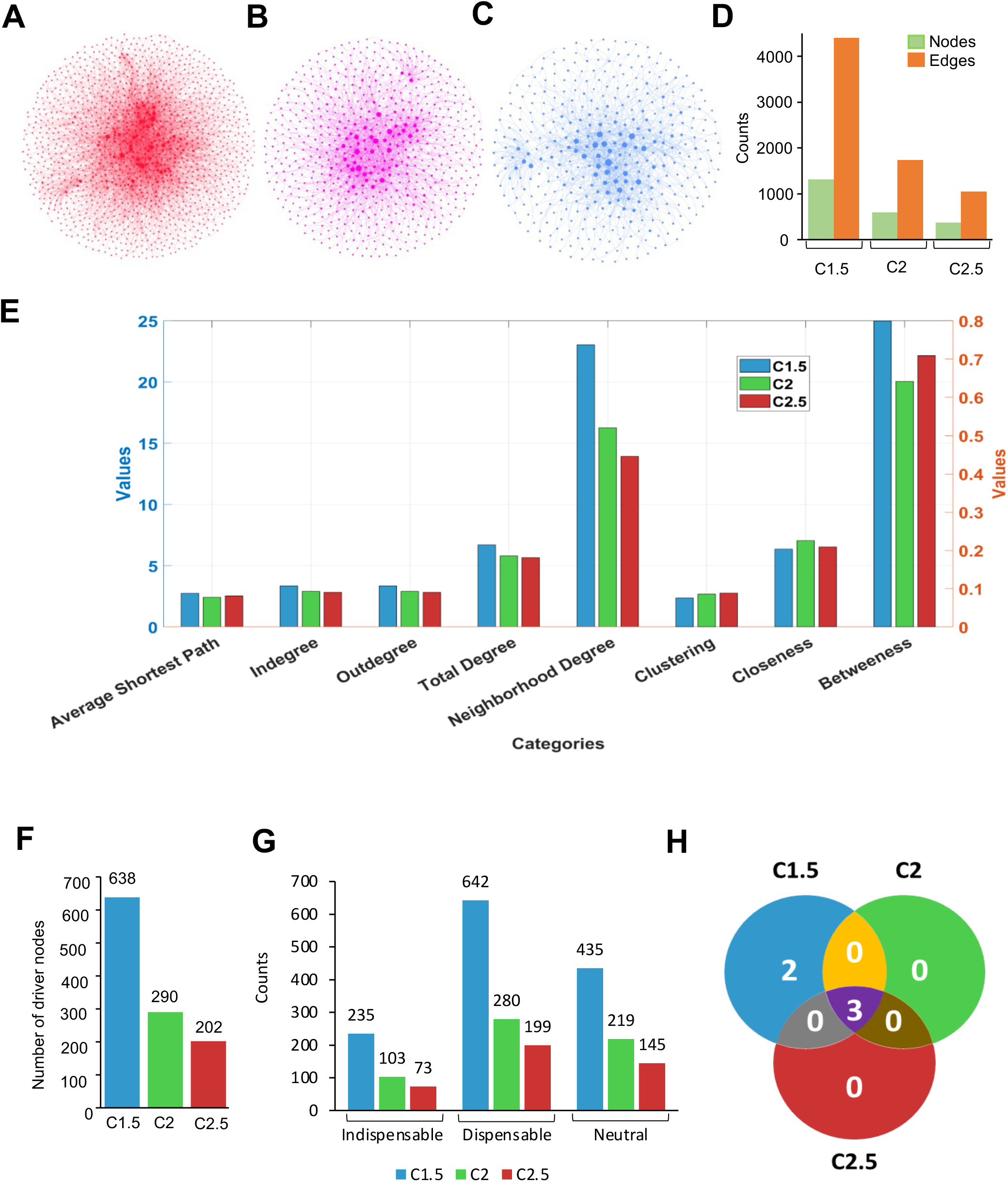
Characteristics of the directional PPI Networks. A-C) DPNs at C1.5, C2, and C2.5 cut off, nodes are sized according to their degree. D) Nodes and degrees in the C1.5, C2, and C2.5 networks. Green and orange bars represent nodes and edges, respectively. E) Topology analysis of the C1.5, C2, and C2.5 networks. Due to variation in the scale, please refer to the right-hand side y-axis (in red colour) for clustering, closeness and betweeness, and for the rest of the categories, please refer to the left-hand side y-axis (in blue colour). F) Driver nodes in the C1.5, C2, and C2.5 networks. G) The number of indispensable, dispensable and neutral nodes in the C1.5, C2, and C2.5 networks. H) Venn diagram showing the potential targets obtained by analysis of the C1.5, C2, and C2.5 networks.

To identify the crucial protein(s) governing this network, we applied the network controllability algorithm (35). The number of driver nodes (minimum number of nodes required to control a network) in C1.5, C2, and C2.5 networks were 638, 290 and 202, which correspond to 49.70%, 48.17%, and 47.72%, respectively (Figure 2F). The large number of driver nodes indicate sparse nature of these networks. Driver nodes were analysed using a simple set theoretic approach to identify indispensable, dispensable, and neutral nodes at each cut off (Figure 2G). Five potential targets (PRKCB IGF1R, DMT1, AKT1 and CSNK1E) were identified at C1.5, of which three (PRKCB, AKT1 and CSNK1E) were retained at C2 and C2.5 (Figure 2H). As per the above analysis, depletion of PRKCB, AKT1 or CSNK1E or functional inhibition of these proteins using biochemical inhibitors should reverse the HEV infection induced change in gene expression profile to that of uninfected cells, thereby acting as an antiviral strategy. PRKCB, AKT1 and CSNK1E encodes PKCβ, AKT1 and CK1ε proteins, respectively.

### Inhibition of g3- and g1-HEV replication by CK1ε and AKT inhibitors

An earlier study reported a lack of any change in HEV replication in cells lacking PKCβ (36). We evaluated the importance of CK1ε and AKT on HEV replication using PF670462 and AKTi-1/2, biochemical inhibitors of CK1δ/ε and AKT1/2, respectively. Huh7 cells were treated with increasing concentrations of PF670462 and AKTi-1/2 (1µM, 5µM, 10µM, 20µM). Twenty four hour treatment with up to 20µM of PF670462 or AKTi-1/2 did not cause any cytotoxicity to Huh7 cells, however 48 hr treatment (2 times treatment at 24 hr interval) with 20µM AKTi-1/2 caused 50% cell death (Figure 3A). Therefore AKTi-1/2 was used at a maximum concentration of 10µM. Huh7 cells expressing *in-vitro* synthesized genomic RNA of g3-HEV replicon (pSK-P6-HEV-Luc) were treated with increasing concentrations of PF670462 and AKTi-1/2 for 24 hr and 48 hr, followed by measurement of *Gaussia*-Luciferase (G-Luc) activity and cell viability. Treatment of g3-HEV-Luc expressing cells with 20µM of PF670462 for 48 hr reduced the luciferase activity by approximately 85% in comparison to untreated cells, while treatment of the same cells with 10µM of AKTi-1/2 for 48 hr reduced luciferase activity by approximately 81% in comparison to untreated cells (Figure 3B).

**Figure 3.**
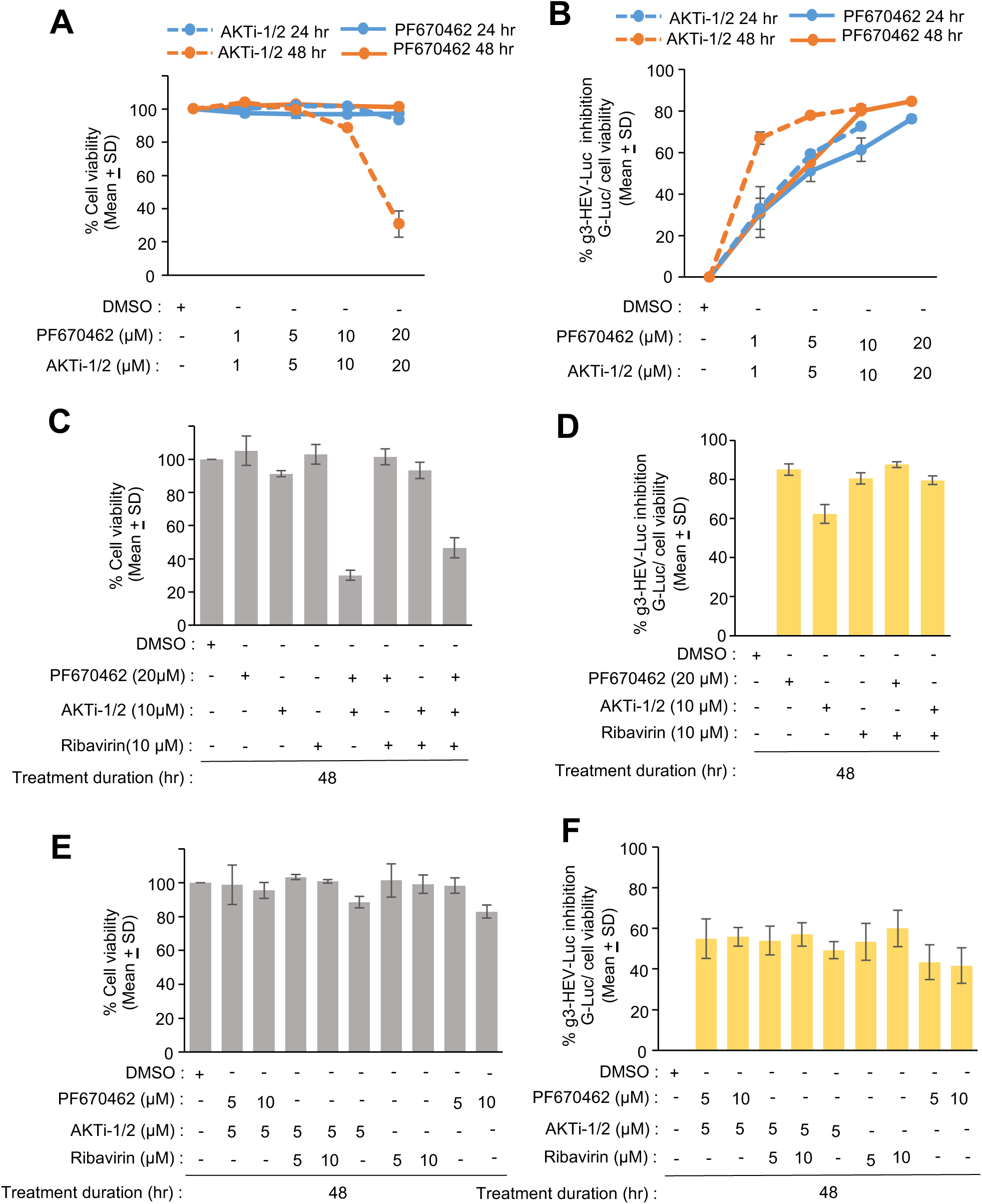
Effect of CK1ε and AKT inhibitors on g3-HEV replicon. A) Percent viability (considering DMSO-treated sample value as 100%) in Huh7 cells expressing P6 HEV-Luc-replicon, treated with indicated concentration of PF670462 (CK1ε inhibitor) and AKTi-1/2 (AKT inhibitor) for 24 hr and 48 hr. Blue and orange colours represent 24 and 48hr inhibitor treatment, respectively. Solid and dashed lines represent cell viability of PF670462 and AKTi-1/2 treated cells, respectively. B) Percent inhibition of *Renilla*-luciferase activity in Huh7 cells mentioned in (A). Solid and dashed lines represent percent g3-HEV inhibition in PF670462 and AKTi-1/2 treated cells, respectively. *Renilla*-luciferase values were normalized to the cell viability values. Percent inhibition was calculated with reference to the values obtained in DMSO-treated samples. Values are mean±SD of three independent experiments. C) Percent viability in Huh7 cells expressing P6 HEV-Luc-replicon and treated with the indicated compounds for 48 hr. Viability of DMSO-treated sample was considered to be 100%, and other values were calculated with reference to that. D) Percent inhibition of *Renilla*-luciferase activity in Huh7 cells mentioned in (C). *Renilla*-luciferase values were normalized to the cell viability values. Percent inhibition was calculated with reference to the values obtained in DMSO-treated samples. Values are mean±SD of three independent experiments. *P* values were calculated using student’s t-test. E) Percent viability in Huh7 cells expressing P6 HEV-Luc-replicon and treated with the indicated compounds for 48 hr. Cell viability of DMSO-treated sample was considered to be 100%, and other values were calculated with reference to that. F) Percent inhibition of *Renilla*-luciferase activity in Huh7 cells mentioned in (E). *Renilla*-luciferase values were normalized to the cell viability values. Percent inhibition was calculated with reference to the values obtained in DMSO-treated samples. Values are mean±SD of three independent experiments. *P* values were calculated using student’s t-test.

Both CK1ε and AKT participate in multiple cellular pathways and act on several targets. It is possible that co-targeting their function might show a cooperative or synergistic or antagonistic effect on inhibiting HEV replication. Ribavirin is an IMPDH (Inosine monophosphate dehydrogenase) inhibitor, which is known to inhibit HEV replication and used as an off-label therapeutic in some HEV cases (7). Possible cooperativity or antagonism between PF670462, AKTi-1/2 and ribavirin on HEV replication was evaluated using the g3-HEV-Luc-replicon model. Huh7 cells expressing the g3-HEV-Luc-replicon were treated for 48 hours with 5, 10 and 20µM PF670462 and 5 or 10µM of AKTi-1/2 or Ribavirin. Cotreatment with 20µM PF670462 and 10µM AKTi-1/2 or 20µM PF670462, 10µM AKTi-1/2 and 10µM ribavirin showed significant cytotoxicity in Huh7 cells (Figure 3C). No significant change in G-Luc activity was observed in g3-HEV-Luc-expressing Huh7 cells upon cotreatment with 20µM PF670462 and 10µM ribavirin or 10µM AKTi-1/2 and 10µM ribavirin (Figure 3D). Further, attempts to optimize the cotreatment dose with varying concentrations of the compounds did not show any cooperative or antagonistic effect (Figure 3E, 3F).

Data obtained in the g3-HEV-replicon model were validated in Huh7 cells expressing g3-HEV infectious virus. Treatment of infectious g3-HEV producing Huh7 cells with PF670462 or AKTi-1/2 significantly reduced the level of viral RNA, in agreement with results obtained in g3-HEV-replicon model (Figure 4A). Immunofluorescent staining of parallelly maintained cells with anti-ORF2 antibody showed a significant reduction in ORF2 positive cells in PF670462 and AKTi-1/2 treated cells (Figure 4B). Quantification of ORF2 positive cells from five representative fields supports the above observation (Figure 4C).

**Figure 4.**
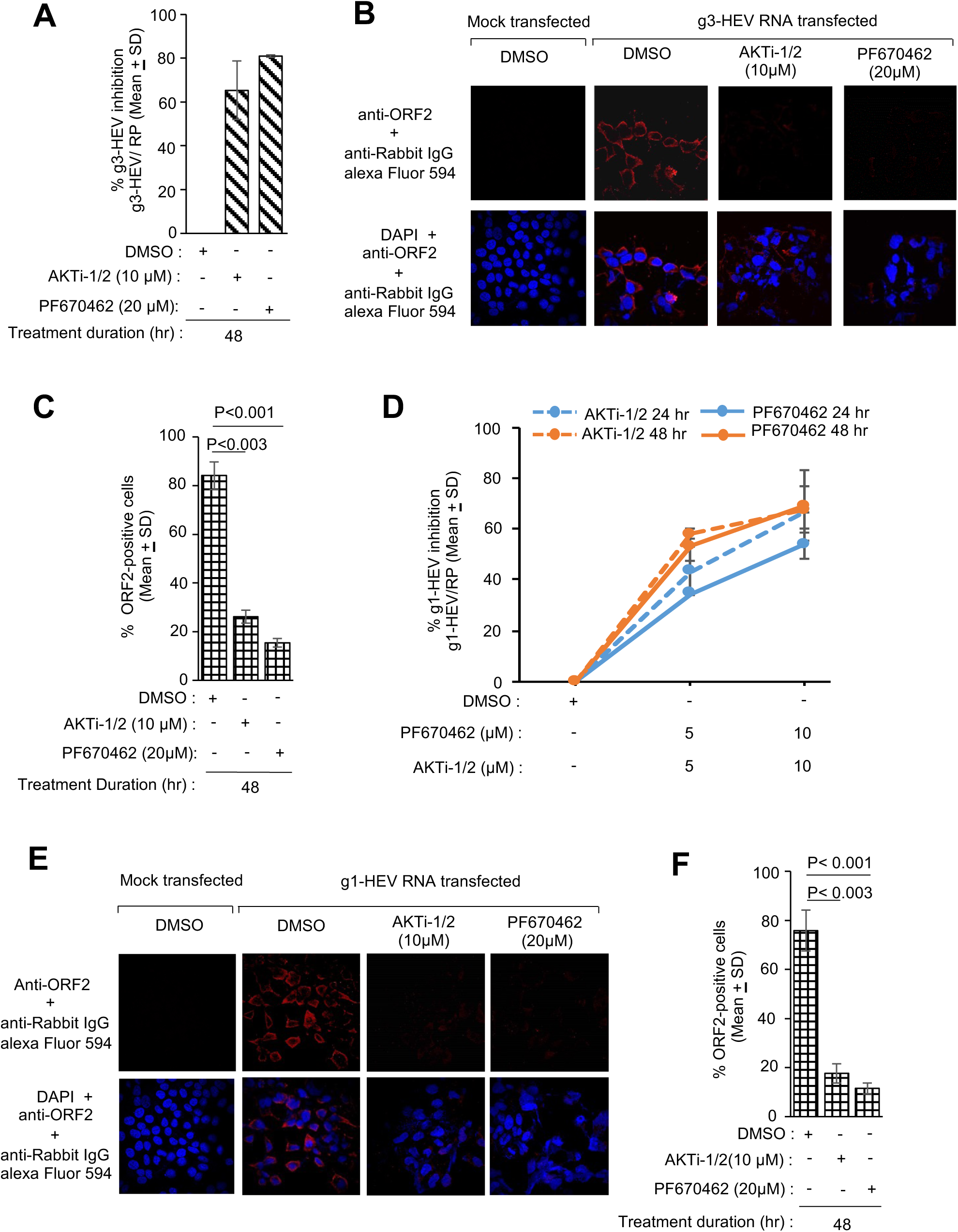
Antiviral activity of CK1ε and AKT inhibitors on g1- and g3-HEV. A) qRT-PCR measurement of percent inhibition of g3-HEV RNA level in Huh7 cells expressing g3-HEV and treated with PF670462 or AKTi-1/2, as indicated. Values are mean±SD of three independent experiments. B) Immunofluorescence image of ORF2 (red, upper panel) or ORF2 (red) and nucleus (Blue) (lower panel) in Huh7 cells expressing g3-HEV and treated with the indicated inhibitors for 48 hours. C) Quantification of % ORF2-positive cells in 5 random fields, as represented in (B). Values are mean±SD of 5 random fields. *P* values were calculated using student t-test. *P*<0.05 was considered significant. D) qRT-PCR measurement of percent inhibition of g1-HEV RNA in Huh7 cells expressing g1-HEV and treated with PF670462 or AKTi-1/2, as indicated. G1-HEV RNA values were normalized with respected to that of RP. Percent inhibition was calculated with reference to the values obtained in DMSO-treated samples. Percent inhibition is plotted as mean±SD of three independent experiments. E) Immunofluorescence image of ORF2 (red, upper panel) or ORF2 (red) and nucleus (Blue) (lower panel) in Huh7 cells expressing g1-HEV and treated with the indicated inhibitors for 48 hours. F) Quantification of % ORF2-positive cells in 5 random fields, as represented in (E). Values are mean±SD of 5 random fields. *P* values were calculated using student t-test. *P*<0.05 was considered significant.

Although all HEV genotypes show a single serotype, differences exist between the genotypes in terms of their host range and interaction with host factors (37, 22). As our initial dataset was derived from g3-HEV infected cells, we evaluated the antiviral activity of PF670462 and AKTi-1/2 on g1-HEV, which is predominant in southeast Asian countries. Huh7 cells were transfected with *in-vitro* synthesized g1-HEV genome (pSK-HEV-2) and treated with increasing concentration of PF670462 (5 µM and 10 µM) and AKTi-1/2 (5 µM and 10 µM), for 24 hr and 48 hr. There was significant reduction in g1-HEV RNA level in 5µM and 10µM PF670462 treated cells at both 24 hr and 48 hr treatment period, later being more efficient (Figure 4D). A similar pattern was seen in AKTi-1/2 treated cells (Figure 4D). Next, an immunofluorescence assay was performed after 48 hr treatment of g1-HEV expressing Huh7 cells with 20µM PF670462 or 10µM AKTi-1/2. As expected, there was a significant reduction in ORF2 signal upon treatment with PF670462 or AKTi-1/2 (Figure 4E, 4F).

### CK1ε is essential for efficient replication of HEV

In order to confirm the role of CK1ε in mediating HEV replication, siRNA against CK1ε was used to deplete the corresponding protein from Huh7 cells, followed by quantification of HEV replication. Huh7 cells were transfected with 25nM of CK1ε siRNA and NT-siRNA (non-targeted siRNA). Seventy two hours post-transfection, viability of the cells were measured by MTS assay. No difference was observed between NT-siRNA and CK1ε siRNA treated cells, ruling out any cytotoxicity of the siRNAs on Huh7 cells (Figure 5A). Western blot analysis of parallelly maintained cells using anti-CK1ε antibody showed marked reduction of CK1ε in corresponding siRNA-treated cells (Figure 5B, upper panel). However, there was no change in CK1δ, (CK1 Delta) protein level, confirming specificity of the siRNA against CK1ε (Figure 5B, middle panel). Level of GAPDH was measured as a control to ensure equal loading of protein (Figure 5B, lower panel). Next, g1-HEV expressing Huh7 cells were treated with the CK1ε siRNA and/or treated with PF670462, followed by quantification of viral RNA level by RT-qPCR. Treatment with CK1ε siRNA or PF670462 inhibited g1-HEV level by ∼70% (Figure 5C). However, cotreatment with siRNA and PF670462 showed no further inhibition (Figure 5C). A similar trend was observed in g3-HEV-Luc expressing cells (Figure 5D).

**Figure 5.**
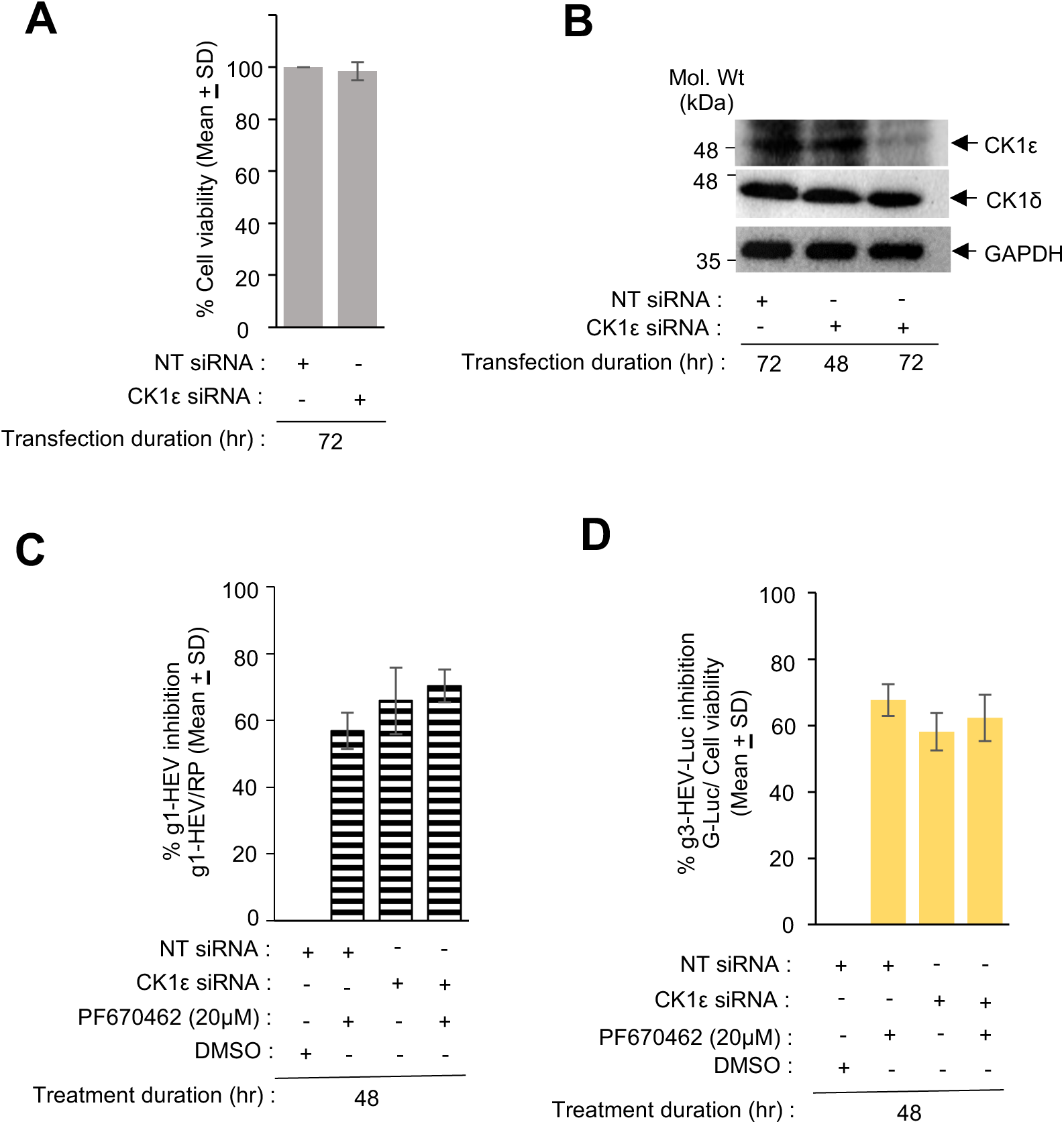
Knockdown of CK1ε inhibits g1- and g3-HEV replication. A) Percent viability in Huh7 cells transfected for 72 hr with NT-siRNA (non-targeting siRNA) or CK1ε siRNA, as indicated. Values of NT-siRNA transfected samples were considered as 100%. B) Western blot analysis of CK1ε (upper panel), CK1δ (middle panel) and GAPDH (lower panel) protein levels in Huh7 cells, transfected with NT-siRNA or CK1ε siRNA for 48 or 72 hr, as indicated. C) qRT-PCR measurement of percent inhibition of g1-HEV RNA in Huh7 cells expressing g1-HEV and transfected with NT-siRNA or CK1ε siRNA for 72 hr and treated with PF670462 or DMSO, as indicated. G1-HEV RNA values were normalized with respected to that of RP. Percent inhibition was calculated with reference to the values obtained in DMSO-treated samples. Percent inhibition is plotted as mean±SD of three independent experiments. D) Percent inhibition of *Renilla*-luciferase activity in Huh7 cells expressing g3-HEV-Luc-replicon and transfected with NT-siRNA or CK1ε siRNA for 72 hr and treated with PF670462 or DMSO, as indicated. *Renilla*-luciferase values were normalized to the cell viability values. Percent inhibition was calculated with reference to the values obtained in DMSO-treated samples. Values are mean±SD of three independent experiments.

### Antiviral activity of Etravirine against HEV

Literature search was performed to identify FDA (Food and Drug Administration, USA) approved drugs that inhibit the activity of CK1ε. Recently, CK1 ε inhibitor Umbralisib was granted FDA approval for the treatment of marginal zone lymphoma and follicular lymphoma (38). However, it was withdrawn following safety issues (39). In another drug repurposing study, FDA approved drug Etravirine was identified as a potential inhibitor of CK1ε (32). Etravirine was originally identified as a non-nucleoside reverse transcriptase inhibitor, used for the treatment of HIV-1 infected patients (28, 29). We evaluated the antiviral activity of Etravirine against HEV.

The possible cytotoxicity effect of Etravirine on Huh7 cells was checked by measuring the viability of Etravirine treated Huh7 cells at 24 hr and 48 hr post treatment with the drug (Figure 6A). Next, the effect of etravirine treatment on g3-HEV replication was measured by using the g3-HEV-Luc-replicon model. Treatment with 10µM Etravirine for 24 hr or 48 hr inhibited virus replication by ∼60% (Figure 6B). Similar inhibitory effect was observed in infectious g3-HEV and infectious g1-HEV expressing Huh7 cells (Figure 6C-6F). IC_50_ (50% inhibitory concentration) of Etravirine against g3-HEV was found to be ∼5µM (Figure 6G). An aliquot of the same samples were tested for cell viability to rule out possible cytotoxicity induced by the drug (Figure 6G). Collectively, these results demonstrate antiviral activity of Etravirine against g1- and g3-HEV.

**Figure 6.**
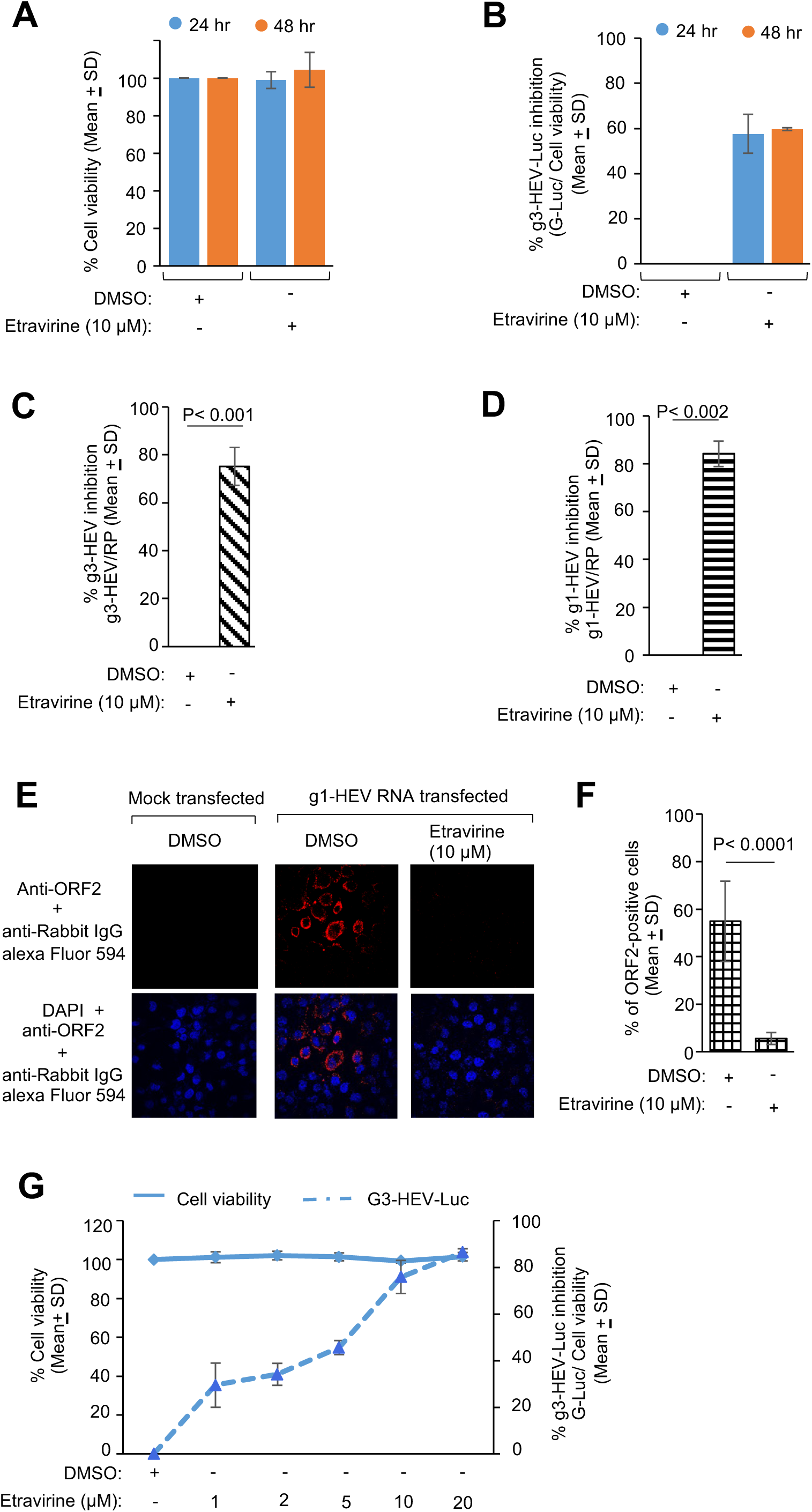
Antiviral activity of Etravirine against g1- and g3-HEV. A) Percent viability in Huh7 cells upon treatment with 10μM Etravirine or DMSO for 24 or 48 hrs, as indicated. Values of DMSO-treated samples were considered as 100%. B) Percent inhibition of *Renilla*-luciferase activity in Huh7 cells expressing g3-HEV-Luc-replicon and treated with 10μM Etravirine or DMSO for 24 or 48 hrs, as indicated. *Renilla*-luciferase values were normalized to the cell viability values. Percent inhibition was calculated with reference to the values obtained in DMSO-treated samples. Values are mean±SD of three independent experiments. C) qRT-PCR measurement of percent inhibition of g3-HEV RNA level in Huh7 cells expressing g3-HEV and treated with 10μM Etravirine for 48 hrs, as indicated. Values are mean±SD of three independent experiments. *P* values were calculated using student t-test. *P*<0.05 was considered significant. D) qRT-PCR measurement of percent inhibition of g1-HEV RNA level in Huh7 cells expressing g1-HEV and treated with 10μM Etravirine for 48 hrs, as indicated. Values are mean±SD of three independent experiments. *P* values were calculated using student t-test. *P*<0.05 was considered significant. E) Immunofluorescence image of ORF2 (red, upper panel) or ORF2 (red) and nucleus (Blue) (lower panel) in Huh7 cells expressing g1-HEV and treated with 10μM Etravirine for 48 hours. F) Quantification of % ORF2-positive cells in 5 random fields, as represented in (E). Values are mean±SD of 5 random fields. *P* values were calculated using student t-test. *P*<0.05 was considered significant. G) Three axis graph showing percent viability (Y axis, left side scale) and percent inhibition of *Renilla*-luciferase activity (Y axis, right side scale) in Huh7 cells treated with 10μM Etravirine or DMSO for 24 hrs, as indicated. Values of DMSO-treated samples were considered as 100%. *Renilla*-luciferase values were normalized to the cell viability values. Percent inhibition was calculated with reference to the values obtained in DMSO-treated samples. Values are mean±SD of three independent experiments. Solid line represents percent cell viability and dashed line represents percent inhibition of g3-HEV-Luc.

## Discussion

In the current study, we followed the computational biology approach to discover novel antiviral targets against HEV and characterized the antiviral potential of AKT/PKB and CK1ε inhibitors. Transcriptome data of HEV infected PHH cells were used as source data for our computational analysis. Compared to transformed cell lines, PHH cells are closer to normal hepatocytes, therefore, the results obtained using PHH-based HEV infection model are likely to be relevant for downstream application. Analysis was started with the identification of DEGs at multiple cut offs, which likely contributes to the overall validity and comprehensiveness of the study’s conclusions. Next, gene reversibility analysis was performed using the CMap database to determine which genes can reverse the differentially expressed gene profile. Next, network controllability analysis was applied to identify the indispensable modulators. Network controllability analysis is one of the most promising algorithms that stands over many of its counterparts (24, 25). It refers to the study and understanding of how to control or influence the behaviour of a complex network by manipulating a small set of strategically chosen nodes. By amalgamating network controllability and disease reversibility, we dissected the regulatory architectures governing gene expression changes following virus propagation and identified the key control sets that shape the emergent behaviours of the disease systems. To improve the robustness of the analysis, we opted for a multiple cut off approach with three different fold-changes of the differentially expressed genes. This approach captures the inherent variability of biological networks that define the regulatory landscape, resulting in a more thorough and nuanced knowledge of the underlying system. By capturing modulators that retain indispensability across a broad range of perturbations, this approach likely increases the potency of the development of broad-spectrum therapeutic strategies with far-reaching clinical implications.

An antiviral against HEV is necessary and attempts have been made in the past in that direction (11–21). However, at present no prescription drug is available against HEV. Literature search did not show any report on anti-HEV activity of PKCβ, AKT1 and CK1ε inhibitors. However, it has been reported that knockdown of PKCβ does not alter HEV replication (36). Therefore, we did not pursue further studies on PKCβ. It is reported that Phosphatidyl inositol three Kinase (PI3K) inhibitor LY294002 and mechanistic target of rapamycin (mTOR) inhibitor rapamycin facilitate HEV replication (40). Similar profile was obtained upon LY294002 treatment of g3-HEV replicon expressing Huh7 cells in our laboratory (data not shown). It is also reported that Phospho-AKT (S473) level is higher in Huh7 cells lacking mTOR as well as cells expressing g3-HEV (40). Upon activation, PI3K stimulates the production of phosphatidylinositol (3,4,5)-trisphosphate (PIP3) PIP3, which binds to the PH-domain of AKT and recruit it to the plasma membrane where 3-Phosphoinositide-dependent kinase 1 (PDK1) phosphorylates Thr308 residue of its kinase domain. Complete activation of AKT requires phosphorylation at its Ser473 residue by any of the following kinases, including PDK-1, integrin linked kinase (ILK), DNA-dependent protein kinase (DNA-PK) or AKT itself. Given the complexity of PI3K-AKT signalling, available data is insufficient to predict proviral or antiviral role of AKT in HEV infected cells.

Activated AKT phosphorylates a variety of downstream proteins such as mTORC1, cell cycle-dependent kinase (CDK) inhibitor p21 and p27, FOXO1, WEE1, GSK3β, IKK (41). Phosphorylation by AKT leads to degradation of FOXO as well as cell cycle inhibitors p21 and p27 (42, 43, 44). AKT mediated phosphorylation also inactivates GSK3, which is a negative regulator of cell cycle progression (45). AKT also suppresses TSC1/2 formation and activates Rheb, an activator of mTORC1, which further phosphorylates ribosomal protein S6 kinase (S6K) and 4E-BP1. Phosphorylation of 4E-BP1 releases eIF4E. Activated S6K and eIF4E promote protein translation and cell proliferation (41).

We investigated the crosstalk between AKT, CK1ε and HEV using respective inhibitors. Both Akti-1/2 and PF670462 (well characterized inhibitors of AKT and CK1δ/ε, respectively) inhibited HEV replication in Huh7 cells. Akti-1/2 is a cell permeable, reversible and selective inhibitor of AKT1/AKT2 activity. Its IC_50_ for AKT1 and AKT2 is 58nM and 210nM, respectively. It binds to the pleckstrin homology (PH) domain of AKT and blocks basal and stimulated phosphorylation/activation of AKT1/AKT2 both in cultured cells *in vitro* and in mice *in vivo* (46). Recently, another AKT inhibitor Capivasertib has been approved for the treatment of adult patients with hormone receptor (HR)-positive, human epidermal growth factor receptor 2 (HER2)-negative localized or metastatic breast cancer (47). Its antiviral potential against HEV should be evaluated.

CK1ε is a serine-threonine kinase of CK1 family. CK1 family has seven members (α, β, γ1, γ2, γ3, δ, ε), in which ck1ε shares highest similarity with ck1δ in the c-terminal domain. CK1ε plays pivotal role in several signaling pathways that control circadian clock, cell proliferation, vesicle trafficking, DNA replication and repair (48,49). Notably, it controls circadian cycle by phosphorylating cytoplasmic PER (period), leading to its degradation; phosphorylates dishevelled 2 (Dvl2), a key component of Wnt/β-catenin pathway, phosphorylates and inactivates 4EBP1, promoting translation initiation (50). Dvl2 phosphorylation by Ck1ε is dependent on its association with DDX3 (considered as a regulatory subunit of Ck1ε) (51). Ck1ε has also been reported to bind to ATP-dependent RNA helicase DDX3X allosterically, activating Wnt signaling (52). On the other hand, AMP-activated Protein kinase (AMPK) phosphorylates Ser389 of Ck1ε and upregulates its kinase activity (53).

Antiviral activity of two Ck1ε inhibitors were evaluated in this study. PF670462 is a selective inhibitor of CK1δ and CK1ε kinases, with IC_50_ of 7.7nm and 14nm, respectively. However, it is not FDA approved. Umbralisib, a dual kinase inhibitor of PI3K-δ and CK1ε was granted FDA approval in 2021 for the treatment of marginal zone lymphoma and follicular lymphoma (38). However, the approval was withdrawn in 2022 due to safety concern (39). Etravirine is a second generation non-nucleoside reverse transcriptase inhibitor (NNRTI). It is FDA approved for the treatment of HIV-1 infected patients. EC_50_ of Etravirine against wild type HIV-1 strain is 4 ng/mL in acutely infected T-cell lines and half-life of Etravirine is ∼41 hours. It is safe for clinical use, minor rash being the side effect in some cases (30). Both inhibitors were equally effective in inhibiting the replication of g1- and g3-HEV. The exact molecular mechanism by which Etravirine and other Ck1ε inhibitors block HEV replication remains to be investigated. We investigated the possibility of HEV RdRp (RNA-dependent RNA polymerase) being a substrate of Ck1ε. Phospho-proteomics analysis of mammalian cell-purified RdRp did not reveal any Ck1ε-phosphorylated residue, thereby ruling out a direct effect of Ck1ε on HEV RdRp (unpublished data). Nonetheless, Ck1ε might be modulating HEV replication by acting on host pathways. For example, DDX3 is known to be a pro-viral factor for HEV replication (54). As mentioned above, DDX3 is a regulatory subunit of Ck1ε. Thus, effect of DDX3 and Ck1ε on HEV replication maybe linked. Moreover, enhanced translation due to CK1ε mediated phosphorylation of 4E-BP1 might facilitate synthesis of viral proteins, which in turn enhances viral replication. Etravirine belongs to the group of diarylpyrimidine derivatives. Recently, additional diarylpyrimidine derivatives were reported to possess better antiviral activity on HIV-1 and lower toxicity, compared to Etravirine (55). Anti-HEV activity of such molecules should be evaluated.

In summary, following computational analysis-mediated identification of antiviral targets against HEV and experimental validation of the targets in mammalian cell culture-based model of HEV replication, this study demonstrates the potent antiviral activity of Etravirine against HEV. Etravirine inhibits both g1- and g3-HEV, which are the most common genotypes to infect humans. Given that etravirine is already used in the treatment of HIV-1 infected patients without any known side effect, it is a promising candidate for further evaluation as an anti-HEV drug in humans. Our study also paves the way for designing novel antiviral strategies against HEV by targeting host Ck1ε and AKT/PKB.

## Materials and Methods

### Computational analysis

The RNA-Seq data reported by Todt et al. was downloaded from GEO database (accession no. GSE135619) (27). Raw data of Control and g3-HEV infected groups was filtered to remove genes with null RPKM in both categories. DEGs were identified at three-fold change cut offs (1.5, 2 and 2.5, Figure 1a). CMap and network controllability analysis was done as described (26). Briefly, the DEGs from different cut offs were independently analysed in CMap database and genes which showed a connectivity score cut-off of −90 or less were selected (56). These genes are hereafter referred to as modulators. A directional protein-protein interaction network (DPN) was generated by combining STRING (version 11.0), Signor (version 2.0) and literature (57). DEGs and modulators of each category were combined and mapped to the DPN to construct three directional subnetworks. Next, structural network controllability analysis was performed following the algorithm published by Liu and co-workers to identify the indispensable nodes in the network (35). Indispensable modulators were identified by a simple set theoretic approach, using the following equation:

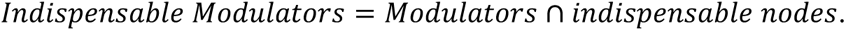

### Plasmids and reagents

The pSK-HEV-2 plasmid (Accession number AF444002.1) and pSK-P6 HEV plasmid (Accession number JQ679013.1), containing the cDNA of g1-HEV and g3-HEV, respectively, have been described (20). The anti-CK1ε antibody (Catalog No. sc-365259) was from Santa Cruz Biotechnology (Texas, USA). The anti-CK1d antibody (Catalog No. 12417) was from Cell Signaling Technology (Massachusetts, USA). The anti-GAPDH antibody (Catalog no. AC001) was from Abclonal (Woburn, MA, USA). The goat anti-Mouse IgG-HRP (Catalog No. 1030-05) was from Southern Biotech (Alabama, USA). mMessage mMachine T7 transcription kit (Catalog No. AM 1344), lipofectamine 3000 (Catalog No. L3000075) and goat anti-Rabbit IgG-HRP (Catalog No. 31460) were from Thermo Fisher Scientific (Massachusetts, USA). PF670462 (Catalog No. S6734) and Etravirine (Catalog No. S3080) were from Selleckchem (Texas, USA). AKTi-1/2 (Catalog No. 124018) was from Sigma (Missouri, USA). DharmaFECT transfection reagent (Catalog No. T-2005) was from Dharmacon (CO,USA). CellTiter 96 aqueous one solution cell proliferation assay kit was from Promega (Madison, USA).

### Mammalian Cell culture, *in vitro* transcription, transfection and cell viability assay

Human hepatoma (Huh7) cells were were maintained in Dulbecco’s modified eagle medium (DMEM) containing 10% fetal bovine serum (FBS), penicillin and streptomycin in a humidified incubator with 5% CO_2_ at 37^0^C, as described (20). pSK-HEV-2 and pSK-P6 HEV plasmids were linearized with Bgl*II* and Mlu*I*, respectively. Linearized plasmids were purified and used for *in vitro* transcription using the mMessage mMachine T7 transcription kit, following the manufacturer’s instruction. g1- and g3-HEV RNA were transfected using lipofectamine 3000 or electroporated into Huh7 cells, as described (20).

For si-RNA mediated gene silencing, Huh7 cells were transfected with 25nM of NT-siRNA or CK1e siRNA ( 5’ GCCGUCGAGAUGACCUGGA 3’), designed using GenScript siRNA target finder tool and synthesized at Eurogentec (Liege, Belgium), using DharmaFECT transfection reagent, as per manufacturer’s recommendation. Stock solution of PF670462, AKTi-1/2, Ribavirin and Etravirine were prepared in DMSO and diluted in culture medium during treatment of the cells. Cell viability was measured using a commercially available kit [CellTiter 96 Aqueous One Solution Cell Proliferation Assay (Promega, Madison, USA)], which utilizes Tetrazolium salt based colorimetric assay, following manufacturer’s instruction.

### RNA isolation, quantitative-Real Time-PCR (qRT-PCR), Luciferase reporter assay, Western blot and Immunofluorescence assay

Total RNA was isolated using TRI reagent (MRC, Massachusetts, USA). qRT-PCR was done using SOLIScript 1-step probe kit according to manufacturer’s recommendation (Solis BioDyne, Tartu, Estonia). Following primers were used: g1-HEV FP: 5’TATACTCGAGGGTGCCGATCGGTCCC3’; g1-HEV RP:5’ TATACCATGGCATCTGGCAGCAAGCTCAG3’; g1-HEV Probe: 5’[FAM] TTGACGCCTGGGAGCGGAATCACC [BHQ1]3’ ; RP FP: 5’AGATTTGGACCTGCGAGCG3’; RP RP: 5’GAGCGGCTGTCTCCACAAGT3’; RP Probe: 5’[FAM]TTCTGACCTGAAGGCTCTGCGCG[BHQ1]3’; g3-HEV FP: 5’TTACGGTTCACGAAGCTCAG3’; g3-HEV RP: 5’GGCGGGTAAGTGCAACTAT3’; g3-HEV probe: 5’[FAM]TGAGACCACAGTTATAGCCACGGC[BHQ1]3’. Gaussia-Luciferase (G-Luc) activity was measured from cell culture media using the *Renilla* luciferase assay kit, following manufacturer’s instruction (Promega, Wisconsin, USA). The G-Luc values were normalized to that of cell viability and plotted as mean (± SD), of three independent experiments done in triplicate. Western blot and immunofluorescence assay was done as described (22).

## Statistical Analysis

Data are presented as mean ± SD of three independent experiments. P-values were calculated by a two-tailed Student t-test (paired two samples for means). A P-value < 0.05 was considered statistically significant.

## Data availability

The RNA-Seq data used in this study is available in GEO database (accession no. GSE135619).

## Acknowledgement

Research in MS laboratory is supported by the THSTI core grant. SA and RV are supported by senior research fellowship from the Council of scientific and industrial research, Government of India. DTS is supported by THSTI intramural project.

## Author Contributions

DTS performed the *in-silico* analyses. SA, RV, KC performed the cell-based experiments. SC supervised the *in-silico* analyses. MS and SC conceptualized, arranged funds and supervised the overall study, all authors participated in writing the manuscript.

## Competing interests

The authors have no interest to declare.

